# Endothelial β-PIX (ARHGEF7) drives exocytosis through enabling the dynamic reorganisation of the cytoskeleton

**DOI:** 10.1101/2025.05.06.652370

**Authors:** Sammy El-Mansi, Karishma Mary Deegan, Lisa Donker, Susana Godinho, Thomas D. Nightingale

## Abstract

Endothelial cells rapidly respond to blood vessel injury or infection by releasing haemostatic and inflammatory proteins from their secretory organelles called Weibel-Palade Bodies (WPBs). Upon stimulation, WPBs traffic to the cell surface to secrete cargo such as von Willebrand factor (VWF), essential for platelet recruitment, and P-selectin, which facilitates the capture and subsequent rolling of immune cells. While this process is critical, many aspects of WPB trafficking and exocytosis remain unclear—particularly how release sites are selected across the cell surface and how fusion is directed to either the apical (lumen-facing) or basolateral (basement membrane-facing) side. These decisions directly impact the biological consequences and effectiveness of the secreted proteins. Using live-cell imaging, fibronectin micropatterning, loss-of-function and biochemical assays, we identified trafficking and release at hot spots near focal adhesions (FAs). We also discovered that the FA-resident guanine nucleotide exchange factor (GEF), β-PIX (ARHGEF7), regulates VWF secretion. β-PIX is known to activate Rho GTPases (Cdc42/Rac), triggering downstream p21-activated kinase 2 (PAK2) signalling and cytoskeletal remodelling. Here, depletion of β-PIX impaired VWF secretion and delayed its release by perturbing cytoskeletal reorganisation. Knockdown and rescue experiments using truncated mutants further revealed which domains of β-PIX are necessary for exocytosis and cytoskeletal reorganisation. This is the first demonstration of β-PIX’s role in VWF secretion from endothelial cells and our data provides new insights into spatial targeting of WPB exocytosis. Such targeting may be essential for guiding leukocyte transmigration or platelet binding, thereby maintaining vascular integrity.

**Schematic:** Endothelial secretory organelles, known as Weibel-Palade bodies (WPBs), store von Willebrand factor (VWF), and their regulated exocytosis is critical for blood clotting and immune responses. This process can be stimulated experimentally by phorbol esters as well as physiological stimuli such as adrenaline and histamine. Cytoskeletal remodelling is required for both pre- and post-fusion events in this secretory pathway. Here, we present evidence supporting the involvement of focal adhesion (FA)-associated exocytosis in the release of WPBs. For the first time, we demonstrate a regulatory role for the FA-resident guanine nucleotide exchange factor (GEF) β-PIX in VWF secretion. Through its GEF activity toward Cdc42 and Rac, β-PIX activates p21-activated kinases, driving the necessary cytoskeletal dynamics. Disruption of β-PIX function results in abnormal cytoskeletal architecture that fails to respond appropriately to phorbol ester stimulation. Consequently, F-actin, nonmuscle myosin IIA (NMIIA), and septins become mislocalised, forming prominent filaments above the nucleus. These data provide a novel insight into the spatial targeting and molecular events underpinning the exocytosis of endothelial secretory organelles.

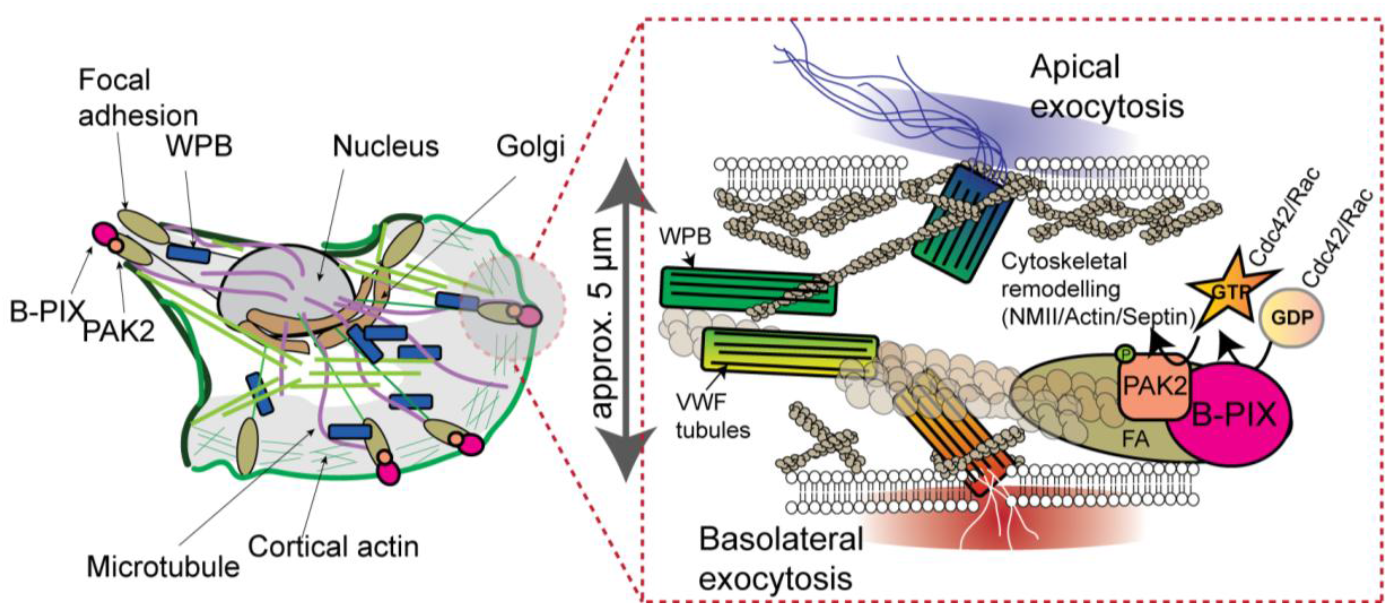

## INTRODUCTION

Weibel Palade Bodies are the regulated storage organelle of endothelial cells. They provide a quickly mobilised pool of proteins that help control inflammation, haemostasis and vascular tone. Contents include, amongst others, the leukocyte adhesion receptor P-selectin and the blood clotting factor von Willebrand factor (VWF).^1,2^ More than 30 different stimuli that cause WPB secretion have been characterised^3^ and these include activators of thrombosis (Thrombin), inflammation (Histamine) and angiogenesis (VEGF). VWF represents more than 90% of WPB content^1,2^ and upon regulated release it unfurls as ultra-long strings into the vascular lumen. This reveals multiple binding sites for circulating platelets and represents the first step of haemostasis. The importance of VWF secretion is best highlighted by patients that release less, or improperly made VWF, that exhibit an extended bleeding pathology termed von Willebrand’s disease.^4^ In addition, it’s clear that VWF has a host of functions beyond haemostasis^5^: during inflammation secreted VWF recruits platelets which helps guide leukocytes to sites of extravasation.^6^ Locally adhered platelets act in a paracrine fashion releasing stored factors (e.g. angiopoetin-1), preventing vascular leakage during leukocyte transmigration^7^.

Spatial regulation of secretion is a fundamental process that underpins cellular function. Examples of targeted secretion include exocytosis at the nerve terminal to allow neurotransmission, release of cytotoxic granules to allow T-cell action and insulin granule release at islet capillaries.^8^ There is evidence of targeted WPB release, VWF secretion from WPB can be either stimulated (for example following exposure to histamine) or basal (a continuous unstimulated release that maintains circulating VWF levels) and both these routes are primarily apical^9,10^. Whereas basolateral VWF secretion is primarily from small constitutive carriers and forms a subendothelial pool. VWF deposition under the cell allows platelet recruitment in the eventuality that endothelial cells are stripped away during injury.^1,9^ Across the breadth of the plasma membrane there is less clear evidence for targeted release. In vitro live cell imaging experiments describe sites of VWF release across many regions of the cell.^11^. This is despite WPB exhibiting a defined maturation process which involves anchoring on peripheral actin structures.^12^

Focal adhesions (FA) are integrin-based, F-actin anchored attachments to the underlying extracellular matrix (ECM). FAs have previously been shown to be trafficking hubs or “hot-spots” for exocytosis of constitutive post Golgi carriers. These carriers transport a variety of cargo that includes matrix metalloproteinases, integrins, collagen, neuropeptide Y and TNF.^13-15^ More recently FA have also been shown to be sites of targeted regulated secretion. Integrin and focal adhesion kinase are activated at the Beta cell capillary interface and this is necessary for glucose dependent insulin secretion^8^.

Although FA targeted release of VWF has not been shown, there is evidence that FA signalling proteins influence VWF exocytosis. Li *et al*., performed an siRNA screen of FA signalling proteins and documented the role of zyxin during cyclic AMP (cAMP)-induced VWF exocytosis. Depletion of zyxin reduced VWF secretion through impaired remodelling of cortical actin and reduced actomyosin ring formation. This corresponded to thrombotic defects *in vivo* by reducing platelet recruitment.^16^

Using a proximity proteomics approach we previously identified that another FA localised protein, p21 Activated Kinase 2 (PAK2), is essential for VWF secretion *in vitro* and *in vivo*^17^. PAK2 catalyses cytoskeletal remodelling and regulates VWF secretion via its *N*-terminal (Cdc42/Rac-binding) activation domain and its *C*-terminal kinase domain.^17^ Our work demonstrated that PAK2 is necessary for the function of a post-fusion actomyosin rings that helps squeeze high molecular weight VWF out of the cell.^11,17-19^ However the physiological importance of its FA localisation to the secretion of VWF secretion is still as yet unclear.

Here we utilised our proximity proteomic data to identify other FA associated proteins with a role in VWF release, this includes the Rho-GEF β-PIX/ARHGEF7. Using micropatterns, biochemical assays and live cell imaging we demonstrate that WPB secretion is in part targeted to FAs and highlight the role of β-PIX in VWF release.

## METHODS

### Cell culture

Human Umbilical Vein Endothelial Cells (HUVEC) (PromoCell, Cat: 12203) from pooled donors were cultured and used as described previously.^20^

### Analysis of proximity proteomics data set

The UniProtKB data base was used to provide a list of known FA associated proteins. The search term “Focal Adhesion Protein” provided a list of proteins that were filtered to obtain those present in *Homo sapiens* with an annotation score of at least 5, these were further restricted to those that were part of the reviewed Swiss-Prot Dataset. This provided a list of 287 protein level entries. We used this list to identify FA associated proteins proximal to WPB from our previously published Rab27a APEX data set^17^. The dataset is publicly available from ProteomeXchange Consortium via the PRIDE partner repository with the dataset identifier 10.6019/PXD036983.

### Fibronectin micropatterning

Coverslips were first cleaned by subsequent incubations in solutions of 100% acetone, 99.9% isopropanol, 99.9% ethanol for 5 minutes each and dried using compressed air to remove any dust. Coverslips were then activated with an ultraviolet (UV) lamp (4Dcell) for 5 minutes. Within 1-2 minutes of removal from the UV machine, the activated side of the coverslips were inverted onto droplets of PLL-PEG (SuSoS) and incubated for 1 hour at room temperature. A bespoke quartz photomask (Toppan) with crossbow micropatterns sized 2500 μm^2^ was then cleaned with 99.9% ethanol, dried with compressed air and activated for 10 minutes with the UV lamp. 5 μl water was added to the activated side of the mask and the PEGylated side of coverslip was dropped on top. The mask was then inverted and placed in the UV lamp such that the UV light passed through the mask onto the side of the coverslip in contact. Coverslips were then detached from the mask by flooding them in water. Dried coverslips were coated with 10ug/ml fibronectin (ThermoFisher) on the patterned side. Fibronectin was washed off with PBS and then a single cell suspension of 7.5 x 10^4^ of cells was seeded on the coverslips in serum free media. After 30 minutes, media was aspirated and replaced with complete media.

### VWF exit site assay

This was performed as described elsewhere^21^ using an anti-VWF (1 in 500 Dako, Cat: A0082) that was conjugated in-house using an Alexa Fluor™ Antibody Labelling Kit (Invitrogen Cat: A20181).

### Immunofluorescence and western blotting

This was performed as described elsewhere^22^ and the commercial suppliers of antibodies used here are provided in Table S1.

### Fixed and live cell imaging

Microscopy studies were performed using either the Zeiss LSM800 or the Nikon CSU-W1 SoRa spinning disk microscope with Z stack intervals ranging from 0.1-0.5 μm depending on the experimental objective. Where necessary image brightness and contrast are adjusted for clarity. 5μg of plasmid DNA was transfected per 0.5 x10^6^ cells. A table detailing the plasmids used for transient transfection are provided in the supplementary materials (Table S2).

### Depth coding of live cell imaging movies

Live cell imaging movies were processed using the Fiji Plugin, Z-stack Depth Color Code and the inversed Rainbow RGB Look Up Table (LUT). Polarity of exocytosis could then be determined by the change in Z depth and colour of expelled VWF.

### siRNA depletion of target proteins in HUVEC

ON-TARGETplus siRNA targeting ARHGEF7 (Cat: L-009616-00-0005), GIT1 (Cat: 020565), GIT2 (CatL-012860-00-0005) and Scribble (Cat: L-010500-00-0005) were purchased from Dharmacon (Horizon). siRNA targeted firefly luciferase transcripts was generated by Eurofins Genomics (sequence 5’ cgu-acg-cgg-aau-acu-ucg 3’). Transfection of HUVEC was performed by electroporation using the HUVEC-old program on the Amaxa Biosystems Nucleofector. Two rounds of 300 pM siRNA were needed to consistently achieve effective knockdown efficiencies (KD) ranging between 70-90% target protein depletion.

### VWF secretion assays

VWF secretion assay and near-infrared (NIR)-fluorescent dot blot were performed as described in our previous research.^23^ Basolateral VWF secretion was determined via this approach through removal of cells using trypsin EDTA and lysing and scraping the underlying ECM in M199 containing 0.5% triton X-100 and protease inhibitor cocktail.

### In vivo

Procedures conducted using mice were in alignment with the institutional Animal Welfare Ethical Review Body (AWERB) and UK Home Office guidelines. Eight-week-old, male, C57BL/6 mice (Charles River, UK) were housed under controlled environmental conditions (12-hour light/dark cycles at ambient temperature and humidity) on a standard chow diet. En face staining of murine aorta was performed as described elsewhere.^24^

## RESULTS

### Focal adhesion associated exocytosis of VWF

To investigate whether FA proteins are localised near WPBs we utilised our Rab27a-GFP-APEX2 proximity proteomic data set.^17^ Using this approach, proteins close to the APEX2 fusion protein resident on WPBs (<20nm) become biotinylated and are purified using streptavidin-coated beads before analysis by mass spectrometry. We compared the FA proteins detected in unstimulated and PMA stimulated endothelial cells and noted a significant enrichment in 78 (unstimulated) and 63 (PMA stimulated) FA proteins including zxyin, vinculin, paxillin and focal adhesion kinase (**Fig. 1A&B**). Using immunofluorescence labelling and confocal microscopy we confirmed that WPB are often localised adjacent to FAs in HUVEC cultures (**Fig. 1C**). We next monitored if WPBs are exocytosed close to FAs (FA-associated exocytosis) by live cell imaging of HUVEC expressing Vinculin-mCherry and GFP-VWF. Upon stimulation with PMA, we observed that WPBs anchored close to FA are secretagogue responsive. Moreover, we could visualise WPBs being trafficked to FA prior to exocytosis (**Fig. 1D and Movie 1**). To determine the proportion of exocytic events that are released apically or basolaterally we utilised Z-stack depth colour coding. Using this approach WPB that fuse and collapse at the basement membrane appear red and generally shift to white upon exocytosis. Whereas apical events are coloured towards the green-blue end of the scale (**Fig. 1E&F, Fig. S1**). VWF secreted into the subendothelial ultimately becomes immobilised and this can be seen below the nucleus where polarity is restricted to release at the basement membrane (**Movie 2**).

**Figure 1:**
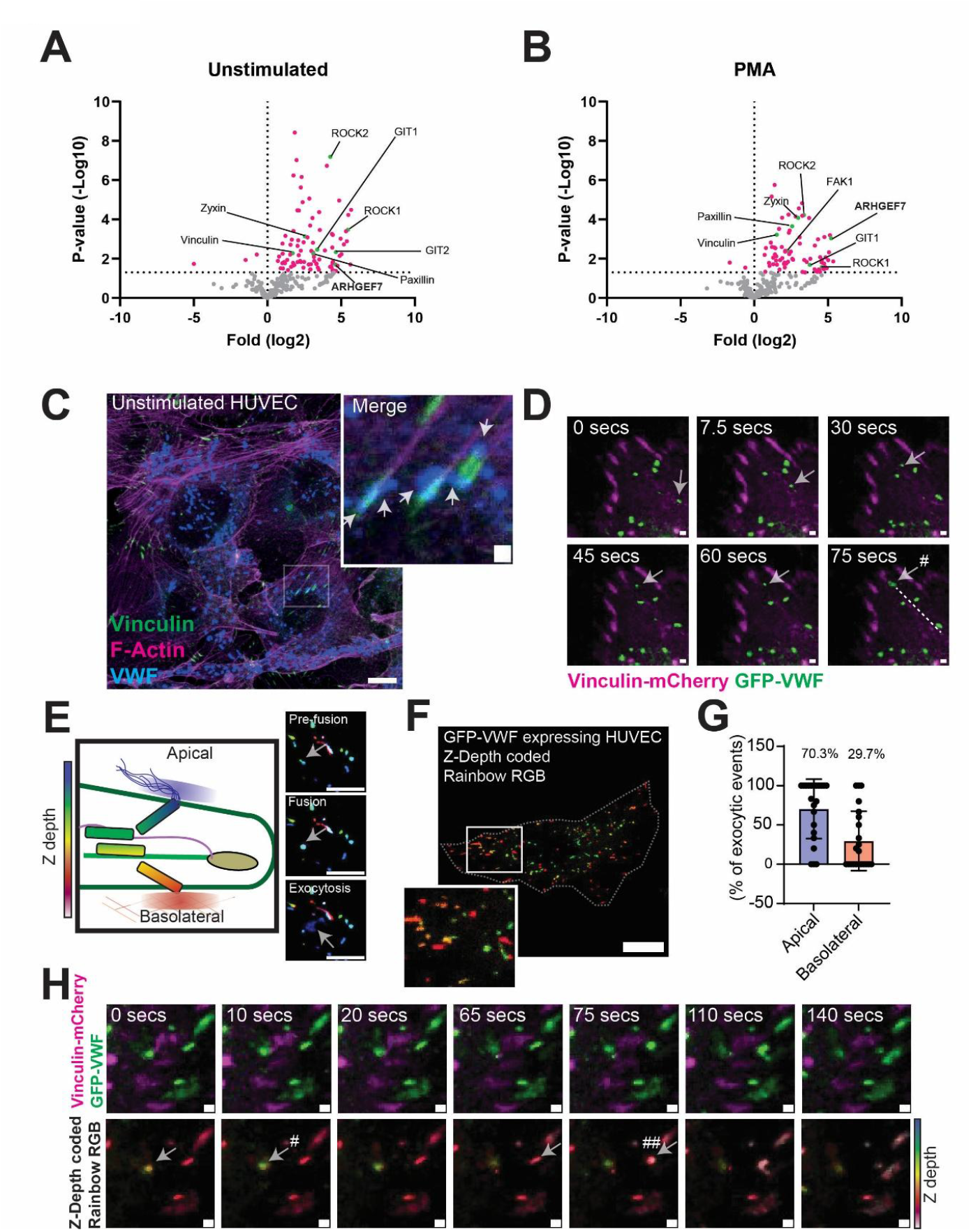
FA-associated WPB exocytosis. **(A)** Volcano plot of FA-associated proteins in close proximity to WPBs, previously identified by Rab27a-targeted APEX2 proximity proteomics (El-Mansi et al., 2023) of unstimulated and **(B)** PMA stimulated HUVEC Grey – not statistically significant (ns) as compared to mock transfected HUVEC. Paired *t* test. Magenta – statistically significant p<0.05, Green – core FA proteins have been highlighted. **(C)** Confocal imaging of vinculin (Green), VWF (Blue) and actin (Magenta). Arrows indicate VWF signal could be seen in rod-shaped structures (WPBs) localized proximal to FAs. Scale bar 10 μm. Inset 1 μm **(D)** Live cell imaging of HUVEC transiently expressing Vinculin-mCherry and GFP-VWF were stimulated with PMA (100ng/mL). Arrow symbol indicates a WPB that is trafficked toward a FA. # indicates point of WPB exocytosis. Dashed line demarks the range of motility across the video. Image is a maximum intensity projection. Scale bar 1 μm **(E)** Schematic of Z depth colour coding confocal live cell imaging movies of GFP-VWF expressing HUVEC to study apical and basolateral polarity. Example stills showing colour change when GFP-VWF is expelled apically. Scale bar 10 μm. **(F)** Additional example of Z-depth colour coding of HUVEC expressing GFP-VWF. Scale bar 10 μm. **(G)** Quantification of the apical-basal polarity of WPB exocytosis in HUVEC stimulated with PMA. (22 cells analysed). **(H)** Image series showing FA-associated WPB exocytosis. Arrows indicate WPB proximal to FAs. # indicates apical exocytosis. ## indicates basolateral exocytosis. Scale bar 1 μm. All videos were acquired through continuous imaging of 0.5 μm interval Z-stacks (5-10 minute movies).

Conversely, signal from apically expelled VWF either dissipated quickly or exhibited increased mobility **(Movie 3**). Following PMA stimulation approximately 70% of events were apical whilst 30% were basolateral (**Fig. 1G**). This closely resembled the distribution of VWF secretion reported through use of transwell cultures.^9^ Applying this approach to live cell imaging movies of HUVEC expressing Vinculin mCherry and GFP-VWF indicated that WPB in close proximity to FAs can exocytosis in either direction (**Fig. 1H**).

Our data indicates that WPB are adjacent to FA and that these represent sites of exocytosis. It also suggests that apical targeting of WPB release is favoured over basolateral release. In vivo research supports these findings as FAs have been observed in large arteries in humans^25^ and mice^26^ and our *En face* imaging of the murine aorta showed VWF signal in alignment with actin structures and near FAs (**Fig. S2**).

Global analysis comparing the localisation of exocytic sites across multiple endothelial cells is difficult due to the heterogeneity of cell shapes. To normalise cell shape and FA distribution we utilised fibronectin micropatterns. If as we hypothesised FA represent hotspots for exocytosis, altering cell shape and sites of adherence should also impact sites of VWF release. Glass cover slips were PEGylated rendering them bio-inert before ultraviolet light exposure through a bespoke quartz mask (Toppan) this allowed us to print crossbow shapes that can be coated in fibronectin/fibrinogen-AF647 (Fig. 2A). We tested three sizes of crossbow pattern before selecting an optimal size of 2500 μm^2^ which provided front-rear polarity and a sufficiently large area to monitor exocytosis (**Fig. 2B**). FAs were prominent along the leading edge and at the tail of the pattern. WPB were observed in close proximity to FAs in both unstimulated (**Fig. 2C**) and stimulated conditions (**Fig. 2D**). To monitor the sites of VWF release more accurately, we stimulated cells in the presence of an Alexa fluor conjugated anti-VWF antibody in the surrounding media (**Fig. 2E**). This allows labelling of VWF as it leaves the cell (exit sites) and effectively distinguished intracellular pools of VWF (WPBs - Blue) from secreted VWF (Red), the latter could be seen in close proximity to vinculin positive FAs (Green) (**Fig. 2F**). Analysis of total VWF suggested that two pools of total VWF exist in the perinuclear and peripheral regions of the cells at approximately equal proportions (54.3±2.8% perinuclear vs 45.7±2.8% peripheral *n*.*s* p>0.05, 56 cells over three independent experiments). In contrast, significantly more external VWF was detected in the cell periphery (23.9±5.1% perinuclear vs 76.1±5.1% peripheral p<0.0001) (**Fig.2 G&H**).

**Figure 2:**
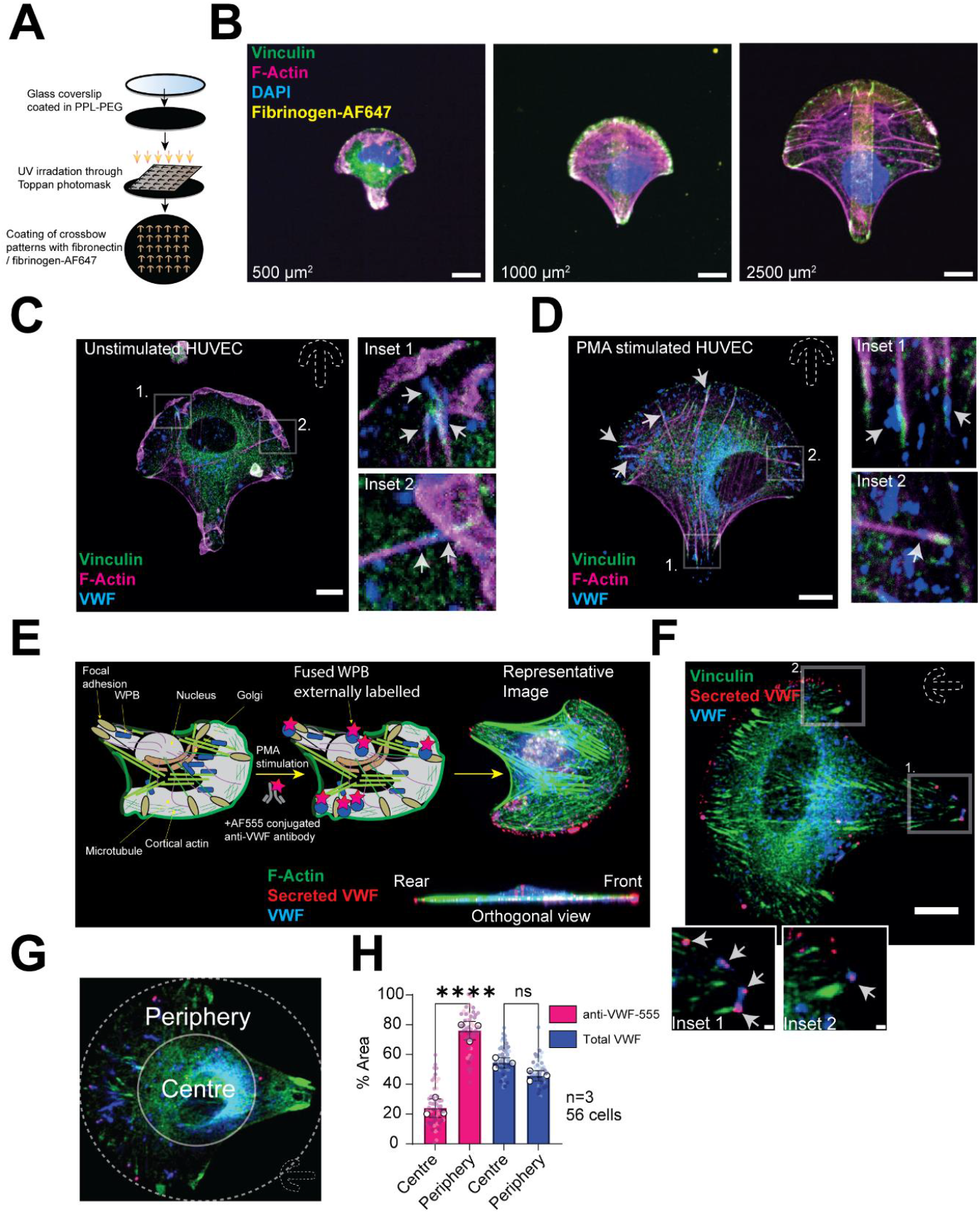
Standardising cell geometry to investigate WPB exocytosis. **(A)** Schematic of process for micropatterning coverslips. A photomask and UV light imprints crossbow micropatterns onto PPL-PEG coated coverslips. Micropatterns can then be coated in extracellular matrix for cell adhesions and standardisation of cell shape and sites of adhesions. **(B)** Fibrinogen-AF647 (yellow) was used to here to visualise and optimise the size of crossbow pattern utilised. Confocal imaging of Vinculin (green), F-actin (Magenta) and DAPI indicated well-defined FAs within 2500 μm^2^ crossbow micropatterns. **(C)** Confocal imaging of unstimulated and **(D)** PMA stimulated cells showed VWF signal (blue) aligning F-actin structures (magenta) and near vinculin-positive FAs (green). Scale bar 10 μm. Arrows show WPB at FAs. **(E)** Schematic of HUVEC on a crossbow micropattern and the strategy to label externalised VWF *in situ*. **(F)** Representative example of extracellular labelling of VWF in stimulated HUVEC. Insets show examples of FA-associated WPB exocytosis. **(G)** Schematic and **(H)** quantification of the proportion of secreted VWF and total VWF in perinuclear and peripheral regions of the cells. N=3 64 cells.

### β-PIX function is required for efficient WPB exocytosis

Using the FA/WPB proximity proteome (**Fig. 1A&B**) we identified candidate machinery that is likely to have a role in targeted WPB exocytosis. We focussed on β-PIX (ARHGEF7) a Rho GEF responsible for activation of Cdc42/Rac and therefore upstream of PAK2 (a kinase which we previously identified is required for VWF release and that is enriched at FAs). Its noteworthy that β-PIX can also directly interact with PAK2 via a PAK-binding SH3 domain. We hypothesised that inhibition of β-PIX may be an alternative approach to limit VWF secretion through targeting of PAK2.

In HUVEC, endogenous β-PIX is localised in the nucleus, cytosol and at one side of FAs (**Fig. 3A**). Overexpression of a mouse Emerald GFP β-PIX Isoform C (Em-β-PIX) and Vinculin-mCherry confirmed this observation (**Fig. 3B**). Using siRNA, β-PIX was effectively depleted (**Fig. 3C**). The amount of regulated VWF secretion in response to PMA was measured using NIR fluorescent dot blot. Depletion of β-PIX prevented apical VWF secretion by 20.1±7.6% cells (Students *t* test p<0.05) (**Fig. 3D**). We also noticed inhibition of secretion using the cell permeable peptide, TAT-PAK18, which disrupts the interaction of PAK and β-PIX (23.3±8.1% decrease p<0.05 Students t test) (**Fig. 3E**).

**Figure 3:**
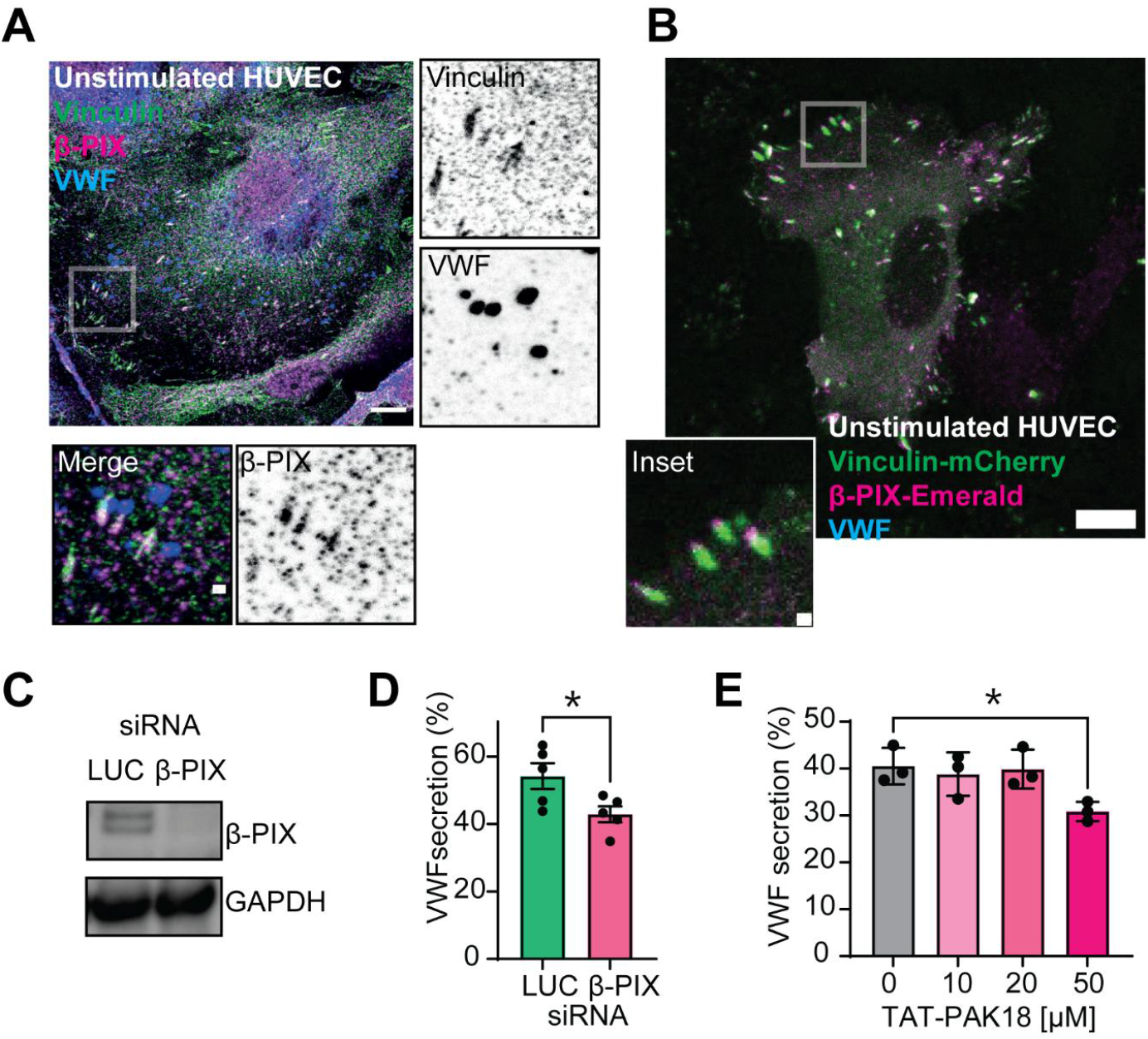
Endothelial β-PIX is resident at FAs and required for VWF exocytosis. **(A)** IF and confocal microscopy of endogenous β-PIX (green), vinculin (magenta) and VWF (blue) shows that β-PIX resides in the cytoplasm, nucleus and FAs. Scale bar 10 um. Insets show WPB in close proximity to FAs. Inset scale bar 1 μm. **(B)** Co-expression of Em-β-PIX and vinculin mCherry shows localisation at one side of the FA. Scale bar 10 μm. Inset 1 μm **(C)** β-PIX was depleted using siRNA and knockdown (KD) efficiency determined by western blotting. **(D)** Luciferase (LUC) control or β-PIX KD HUVEC were exposed to PMA (100 ng/mL) for 30 minutes before the amount of VWF secreted into the culture media was quantified using Near-infrared fluorescent dot blot. N=5 **(E)** Disruption of β-PIX - PAK2 interaction through preincubation with the cell permeable peptide inhibitor TAT-PAK18 significantly reduced PMA induced VWF secretion at the highest concentration of 50 μM. *p<0.05 One way ANOVA with Dunnett’s multiple comparisons test. N=3.

Through western immunoblotting we assayed the activation status of PAK2 in the presence or absence of siRNA targeting β-PIX and monitored phosphorylation at threonine 402. PAK2 activation was dependent on β-PIX (**Fig. S3**). Supporting this, phosphorylation of cofilin (an actin severing protein downstream of PAK2) was significantly reduced in β-PIX KD cells (**Fig. S4**). Other PMA-mediated intracellular phosphorylation events (e.g. ERK1/2) were unaffected. Taken together these data suggest that β-PIX is required for VWF release and is upstream of Cdc42/Rac/PAK2 with a direct role for the interaction of β-PIX and PAK2.

### Depletion of β-PIX alters the dynamics of WPB exocytosis

We next performed quantitative analysis of WPB fusion events, utilising live cell imaging of HUVEC expressing GFP-VWF and a soluble fusion marker, P.selectin luminal domain mCherry (P.sel.lum.mCherry) (**Fig. 4A**). PMA stimulation led to ∼50% reduction in WPB fusion over period of 5 minutes (Students *t* test p<0.05) (**Fig. 4B**). The time delay (lag time) between loss of P.sel.lum.mCherry and loss of GFP-VWF following fusion and collapse of the WPB indicates the efficiency of VWF release post-fusion. WPBs recruit actomyosin rings to aid this process; therefore, inhibitors of actin and myosin delay GFP-VWF release post-fusion.^11,18^ Here, β-PIX depletion significantly extended the time between loss of P.sel.lum.mCherry and the disappearance of spherical GFP-VWF signal (27±5 secs vs 101.3±4 secs, p<0.0001) (**Fig. 4C**).

**Figure 4:**
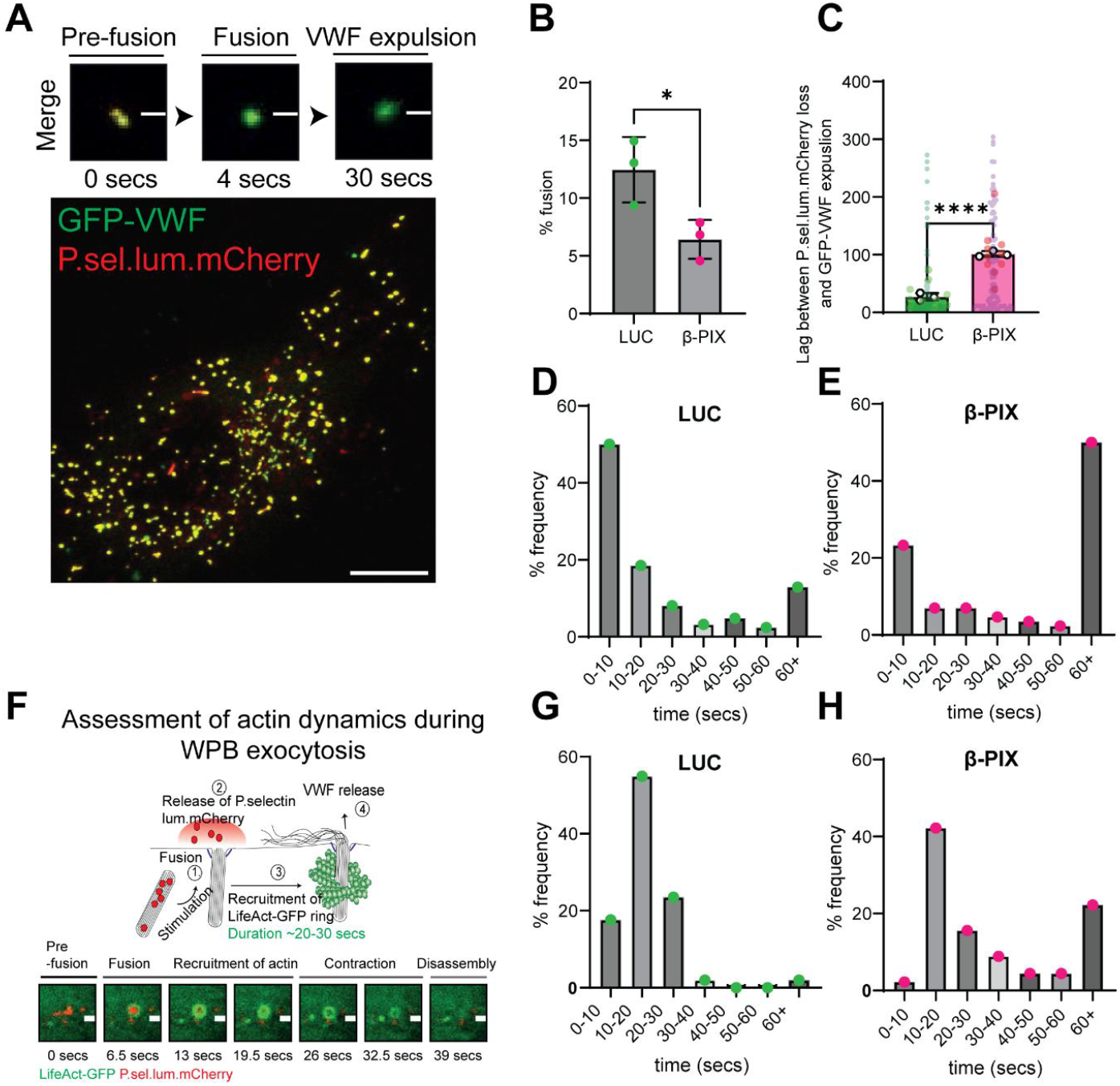
Depletion of endothelial β-PIX influences WPB exocytosis through impeding cytoskeletal remodelling. **(A)** Live cell imaging was performed to assess WPB fusion in control and β-PIX KD HUVEC. Example of HUVEC transiently expressing GFP-VWF (green) and P.sel.lum.mCherry (red). Scale bar 10 μm. **(B)** Mean percentage of WPB fusion events within 5 minutes. 22 cells analysed over three independent experiments. Students *t* test. P<0.05 **(C)** Mean lag time between loss of fusion marker and GFP-VWF expulsion. 208 events analysed in 22 cells over three independent experiments. Students *t* test. P<0.001 **(D)** Distribution of frequency of events in Control (LUC) and **(E)** β-PIX KD cells. **(F)** Schematic for the assessment of actin dynamics during WPB exocytosis. **(G)** Distribution of frequency in the duration of LifeAct-GFP actin rings in **(G)** LUC and **(H)** β-PIX (96 events in 25 cells over three experiments).

This was reflected by a shift in the distributions of frequencies of events lasting less than 10 seconds and more than 60 seconds (**Fig. 4D&E**). Analyses of actin dynamics during WPB exocytosis (**Fig. 4F**) determined that β-PIX KD resulted in a shift in the proportion of events where actin rings persisted for longer. This is indicative of inefficient release (**Fig. 4G&H**).

### Regulated β-PIX-mediated cytoskeletal remodelling requires PAK and Cdc42/Rac binding domains

β-PIX forms subcomplexes with G-protein-coupled Receptor Kinase Interactors (GIT1/2)^27^ and in certain cells Scribble^28^ these interactions are thought to be required for β-PIX function. Using siRNA depletion and NIR blotting we found no evidence for a role of either of these proteins in VWF secretion (**Fig. S5**) To further understand how β-PIX is localised and how it controls VWF secretion and cytoskeletal modelling we used Emerald-GFP tagged β-PIX truncated mutants^29^ (Em-β-PIX) (**Fig. 5A**).

**Figure 5:**
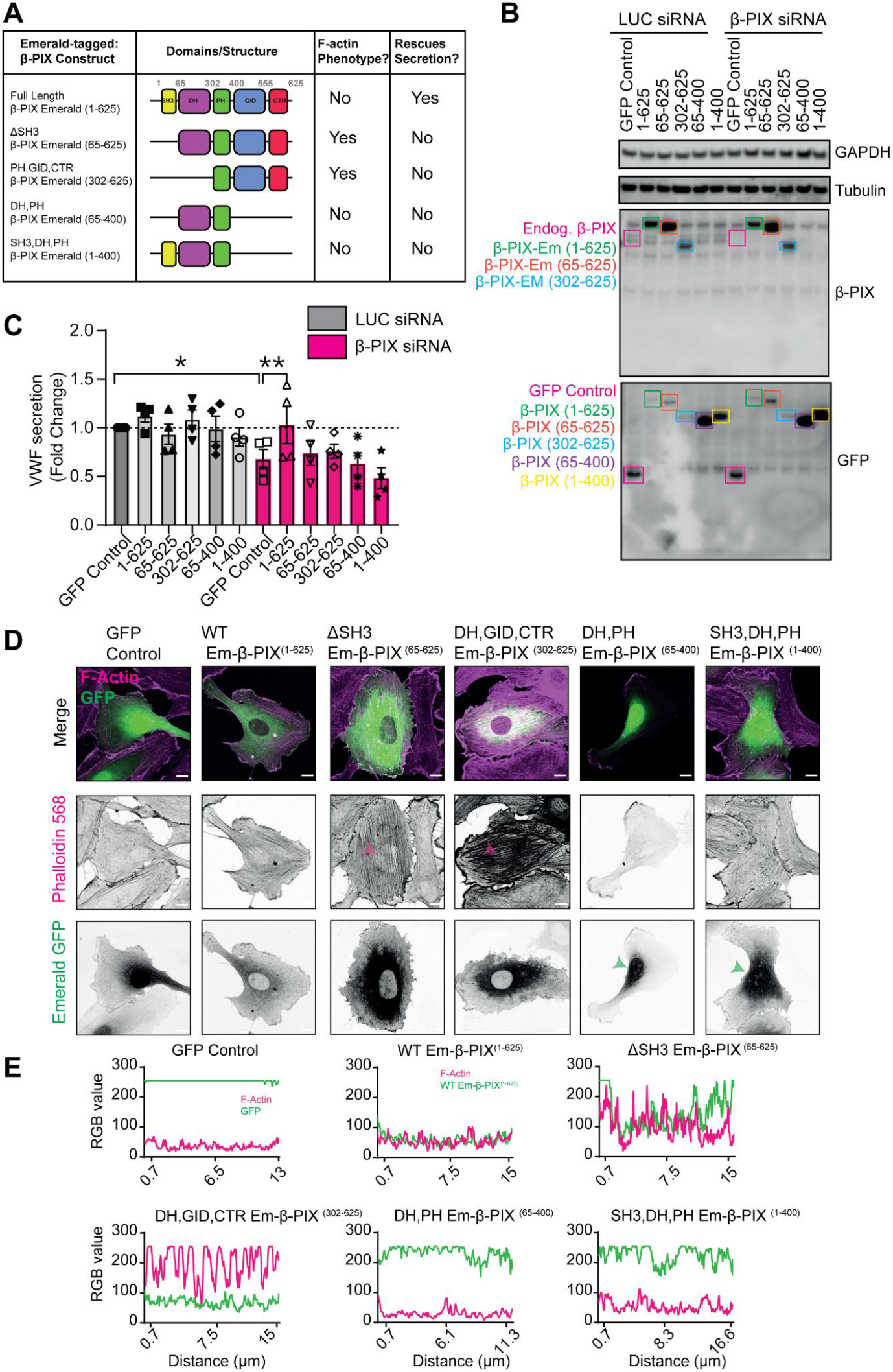
β-PIX requires all of its functional domains for VWF secretion. **(A)** Schematic representation of the domains and structure of Emerald-GFP β-PIX (Em-β-PIX) constructs and their effect on the cytoskeletal architecture. **(B)** Knock down and rescue experiments were performed by transfecting LUC and β-PIX KD HUVEC with 5μg plasmid construct before assaying 24 hours later. Western blotting was used to detect target protein depletion (Magenta box - endogenous β-PIX) and construct expression using anti-β-PIX. A GFP antibody was used to detect the constructs lacking the C-terminal epitope (Purple and Yellow boxes). **(C)** KD and rescue HUVEC were stimulated with PMA (100 ng/mL) and VWF secretion quantified through NIR dot blot. Only the full length Em-β-PIX construct restored PMA induced VWF secretion in β-PIX depleted cells. Data is presented as a fold change. Ratio paired t test. *p<0.05 **p<0.01. **(D)** Overexpression of truncated mutants have dominant negative effects on cytoskeletal architecture. Confocal images of F-actin structure (magenta) and Em-β-PIX (green) localisation. Green arrows indicate mutant constructs lacking the GID and CTR translocate to the nucleus. Magenta arrows indicate dense F-Actin dorsal stress fibres. Note differences in neighbouring cells which do not express the construct. Scale bars are 10 μm. **(E)** Localisation of Em-β-PIX and the F-actin architecture was assessed by plotting the RGB profile above the nucleus.

β-PIX has five domains, an *N*-terminal SH3 domain (PAK-binding), a Dbl-homology domain (DH) with Rac1 and Cdc42 GEF activity, a lipid binding plekstrin homology domain (PH), a GIT1/2 Interacting Domain (GID), and a poorly understood *C*-terminal Region (CTR). We performed a knock down and rescue experiment to assess the effect of each domain on VWF secretion. Western immunoblotting determined efficient KD of endogenous β-PIX (Magenta boxes) and expression of the full-length Em-β-PIX (1-625) as well as the truncated mutants in both Luciferase (LUC) and β-PIX KD cells (**Fig. 5B**). The β-PIX antibody used was raised against a *C*-terminal located epitope, hence anti-GFP antibodies were required to detect Em-β-PIX (65-400) and (1-400). Only overexpression of the full-length Em-β-PIX was sufficient to rescue the defect in regulated VWF secretion (Ratio paired *t* test, *p<0.05). Indicating that multiple binding interactions mediated by the various domains are required.

We also assessed the effect of the various mutant constructs on the endothelial cell cytoskeleton. We noticed two cytoskeletal phenotypes (**Fig. 5D**). Firstly, truncation of the GID and CTR resulted in nuclear localisation of Em-β-PIX^(1-400&65-400)^ (Green arrows). Secondly overexpression of mutant Em-β-PIX constructs lacking the SH3 or DH domain phenocopied the effect of β-PIX depletion on the cytoskeletal architecture (Magenta arrows: dense F-Actin dorsal stress fibres) (**Fig. 5D**). HUVEC expressing Em-β-PIX^(1-400)^ or Em-β-PIX^(65-400)^ did not show dominant negative effects in this regard. RGB profile plots of GFP and the actin cytoskeleton confirm these phenotypes (**Fig. 5E**).

To further understand the importance of the Cdc42/Rac activating GEF domain we utilised a GEF-dead point mutant Em-β-PIX^(1-625 L238R/L239S)^ (**Fig. S6A**). This phenocopied the effect of delta SH3 or SH3/DH mutants (**Fig. S6B&C**). Moreover, IF of endogenous non-muscle myosin II (NMII) demonstrated its misplacement in Em-β-PIX (1-625 L238R/L239S) transfected cells (**Fig. S6D&E**).

This was confirmed with a GFP-tagged NMIIA construct. (**Fig. S7**). Moreover, we noted these dense fibres were rich in Septin-7. This phenotype was rescued through the overexpression of a constitutively active Cdc42 (Cdc42-GFP-V12) fusion protein (**Fig. S8**). Interestingly, Cdc42-GFP-V12 expression also resulted in the formation of rings positive for Cdc42-GFP-V12, actin and septins. Cdc42-V12 activates PAK2 and this supports our previous findings that implicated PAK2 in septin ring formation post-fusion.

The difference between the secretory and cytoskeletal phenotypes indicate that the role of β-PIX in secretion is not dependent on one aspect of cytoskeletal remodelling. More likely that it is the coordination of this and an interaction with other FA proteins that is required for function.

## DISCUSSION

Our work provides new evidence in primary endothelial cells that secretion of WPBs is targeted. With an increased likelihood of release occurring at, or near, FAs. Such localisation is difficult to characterise in a diverse monolayer but we demonstrate that controlling cellular shape using fibronectin micropatterns makes such release apparent. FAs have been characterised as sites of release for constitutive carriers for some time, with MMP trafficking providing a means to remodel FAs^13^ and a more general role in the transport of Rab6 carriers containing TNF, VSVG, gp135 amongst others.^15^ Notably, in the lymphatic endothelium Rab6+ vesicles containing chemokines (CCL21) are spatially targeted to multicellular junctions to promote dendritic cell transmigration.^30^

To our knowledge there is only one example of another regulatory released organelle being preferentially targeted to FAs and this is during insulin granule secretion. This focal release allows secretion of insulin granules into capillaries. Granule fusion targeting is abolished in a β-cell glucotoxicity model suggesting that aberrant targeting may be a component of disease pathology.^31^ We are just beginning to understand the reason for focal release of WPBs and its relevance in vivo, it could expedite location of specific biological effects such as immune cell interaction with the blood vessel.

Alternatively delivery of VWF in vivo is potentially more targeted to FA junctions (FAJ) which are a more common feature.^32^ There are described roles for VWF and platelet capture in controlling vascular leakage and delivery to the cell junctions would favour this.^7^

It is thought that regulated VWF release is primarily apically polarised^9^ and given sites of adhesion are basolateral (at least in culture) this would appear inconsistent with it being a site of targeted release. We utilised a three-dimensional live cell imaging approach and by colour coding different cellular depths, we could accurately track whether VWF was secreted basolaterally or apically. Our findings are consistent with previous reports^9^ with approximately 70% of WPB body fusion occurs on the apical surface. By imaging we noted WPB adjacent to FA exocytosing in both the apical and basolateral direction, demonstrating that both routes are possible from the same FA site. This is perhaps not surprising when you consider the endothelium can measure as little as 0.1 μm at its thinnest point (and a maximum of 10 μm at the nucleus) whilst by comparison WPB are 0.2-0.3 μm wide and as long as 5μm, easily reaching across the depth of a cell in most regions.^33^ 95% of WPBs fuse at just one end, a phenomenon known as “tip-end fusion”.^34^ This indicates that pools of WPB that are at the periphery of the cell at, or near, FA can by shifting on their axis be released at either the apical or basolateral aspect. The only exception being, where a WPB is positioned above or below the nucleus, in these situations only one membrane is accessible and this is corroborated by our live cell imaging.

Physiologically apical or basolateral release of WPB is causative for different functions with the impact varying depending on the cargo under consideration. For VWF, string formation and platelet capture occurs at the apical side whereas basolateral secretion is necessary to generate a subendothelial pool of VWF in case the endothelial surface is stripped during vascular damage. P-selectin is a leukocyte capture receptor and only apical secretion would deliver the cargo to the relevant aspect of the cell. Whilst other cargo including cytokines (IL-8, IL-6 and RANTES), angiogenic molecules (Ang2/IGFBP7) and tonicity modulators (Endothelin) could potentially have effects whatever the polarity of secretion.

Different WPB mobilising stimuli have the potential to alter the likelihood of apical or basolateral release further tuning the physiological response and this might vary in a pathological setting. It will be important to determine if and how polarity of exocytosis is controlled with these alternate stimuli and their dependence or independence of FA release.

During pathological vascular remodelling such as atherosclerosis, endothelial cells are inflamed and VWF is commonly observed in the neointima of blood vessels. Such deposition is exclusively derived from the endothelium^35^ and indicates that endothelial dysfunction can increase basolateral secretion of VWF. As VWF promotes vascular smooth muscle cell proliferation and migration through integrin signalling, misdirected VWF release into the neointima has the potential to contribute to the adverse vascular remodelling observed in in pulmonary hypertension and atherosclerosis.^36^ Furthermore, inflamed endothelial cells upregulate gene expression of cytokines such as IL-6 and IL-8 which are then co-stored in WPBs. Basolateral secretion of such proinflammatory cargo could be having profound effects in the intima of large blood vessels under pathophysiological conditions.

Our mass proximity mass spectrometry demonstrated that approximately 65 FA proteins are enriched at or near WPB both before and after stimulus. Using biochemical assays and live cell imaging we showed that β-PIX facilitates VWF secretion at the apical surface of ECs. Depletion of β-PIX reduced the proportion of WPBs that fused with the PM. Furthermore, the time taken for GFP-VWF to be expelled, following the fusion and collapse of the granule was extended illustrating both pre- and post-fusion roles. We also noted prolonged assembly and contraction kinetics of exocytic actomyosin rings/coats which form post-fusion and aid the expulsion of ultra large VWF multimers.^11^ Previously zyxin another FA localised protein has been implicated in the release of VWF by controlling the formation and maintenance of cortical actin remodelling following cAMP stimulation^16^. Together this suggests that near FAs a region of dynamic actin remodelling and nucleation exists that favours VWF expulsion perhaps by enriching certain key pieces of machinery.

We further interrogated the domain structure of β-PIX to define functional domains. We noted distinct dominant negative phenotypes on actin structures of the various mutants. Prominent F-actin stress fibres were apparent when the PAK-interacting SH3 domains and Cdc42/Rac interacting DH (GEF) domain were included and point mutations in the DH were sufficient to recapitulate this phenotype.

This suggests that β-PIX acts to corral PAK2 via the SH3 domain whereas the GEF domain provides a means to instigate this signalling cascade through activation of Cdc42/Rac. Depletion of β-PIX results in prominent F-actin stress fibres above the nucleus that are decorated with NMII and septins. This cytoskeletal architecture was not affected by PMA stimulation, however, could be altered through expression of a constitutively active GFP-tagged Cdc42 construct (V12). Notably a similar phenotype has been observed in cancer associated fibroblasts and MDCK cells.^37,38^ When monitoring VWF secretion only the full-length Em-β-PIX was able to restore the defect caused by the depletion of endogenous β-PIX. Surprisingly, constructs lacking the GID (GIT1 interacting) and CTR domains failed to rescue the phenotype, despite retaining the ability to bind PAK and Cdc42/Rac. We propose that this failure is due to altered intracellular localization, as these constructs were predominantly sequestered in the nucleus.

From our work it appears that β-PIX functions efficiently as an actin bound or cytosolic pool when its scaffolding proteins are depleted. Scribble localises β-PIX both to the junctions and PM^28^ whereas GIT1 recruits β-PIX to FAs.^27,39^ Depletion of Scribble and GIT1/2 did not effect on VWF secretion. Our data suggests either that Scribble or GIT isoforms have redundant roles and compensate for each other or that β-PIX is recruited to FA by additional means. We hypothesise that under control conditions, PAK2 is recruited to FAs through β-PIX^40^ allowing for spatially restricted control of signalling and cytoskeletal remodelling.^39^

We noted an enriched pool of WPB at FAs which likely reflects direct transport to these sites. WPB are transported on microtubules (MTs)^41^ and we previously demonstrated that the plus end motor protein KIF5B is enriched on one end of WPB.^17^ Delivery to sites of FAs in insulin secreting,^42^ non-neuronal cells such as keratinocytes^13^ and HeLa cells^15^ relies on so called active zone proteins (LL5β, liprin, ELKS and KANK) which form FA associated complexes which mediate CLASP mediated attachment of MTs. From our WPB proximity data set we have noted enrichment of some of these proteins at WPB (CLASP1,2, Kank3, Liprin, LL5β) but are yet to investigate if they are sufficient to target WPBs and MTs to FAs. Previous work has demonstrated that cAMP mediated secretagogues and to some extent Histamine mediate dynein dependent perinuclear clustering of certain WPB preventing their release.^3,43^ Such transport mediated by Rab46 may represent a way of selecting and controlling the release of alternate “pools” of WPB that contain specific cargo^43^. Although how the alternate pools are generated and sensed by Rab46 remains to be determined. There remains the possibility that discrete trafficking machinery is required for specific secretory events e.g. apical or basolateral WPB release.

The in vivo importance of FA targeted release in blood vessels is yet to be determined. Surprisingly, the mere existence of FAs *in vivo* has been a topic of debate within the community. This controversy was addressed by Van Geeman *et al*., who conducted *en face* imaging of human mammary arteries and umbilical veins.^32^ Well-defined, F-actin anchored, FAs were only observed in the arterial vessel at cell boundaries, but not in the vein. This might indicate that sites of WPB release might similarly vary in artery vs vein. The authors proposed the disparity might stem from the differences in biomechanical properties experienced in the venous or arterial system including changes in shear and ECM. As blood vessels age, their biomechanical properties change, becoming stiffer, it would be interesting to determine how this might change FA structure, function and secretion in a living organism. It may not be coincidental that arterial vessels, respond more robustly to the systemic administration of adrenaline (∼80% decrease in endothelial stored VWF over 30 minutes).^44^ This might indicate that the FA profile of vessels and their underlying ECM stiffness plays a specialised role in mediating their responsiveness to physiological stimuli. This also hints to a link between biomechanical adaption during aging and increased haemostasis. Future work should address this knowledge gap in the field.

In summary, we have identified WPB release occurs at hotspots near FA and a novel role for β-PIX in the scaffolding and activation of PAK2 which aids VWF secretion through control of cytoskeletal architecture. How this targeting varies in different vascular beds and in physiological and pathological scenarios will be an important area of future investigation.

## ACKNOWLEDGMENTS

The authors thank their funders and collaborators. This work was supported by the British Heart Foundation (Grant PG/22/11208) and Queen Mary University of London (PhD Studentship: PRN1040R). We acknowledge the CMR Advanced Bio-Imaging Facility of QMUL for the help and advice with microscopy.

## AUTHORSHIP CONTRIBUTIONS

S.E.-M and T.D.N developed the methodology.

S.E.-M and K.M.D performed the investigation.

L.D optimised and trained the investigators in fibronectin micropatterning techniques.

S.G provided valuable tools and guidance.

S.E.-M. and T.D.N wrote the original draft; and all authors reviewed and edited the manuscript.

T.D.N. supervised the study.

## AUTHORSHIP CONTRIBUTIONS DISCLOSURE OF CONFLICTS OF INTEREST

The authors declare no competing financial interests.

## REFERENCES

1. McCormack JJ, Lopes da Silva M, Ferraro F, Patella F, Cutler DF. Weibel-Palade bodies at a glance. Journal of cell science. Nov 1 2017;130(21):3611–3617. doi:10.1242/jcs.208033

2. Hordijk S, Carter T, Bierings R. A new look at an old body: molecular determinants of Weibel-Palade body composition and von Willebrand factor exocytosis. J Thromb Haemost. May 2024;22(5):1290–1303. doi:10.1016/j.jtha.2024.01.015

3. Rondaij MG, Bierings R, Kragt A, et al. Dynein-dynactin complex mediates protein kinase A-dependent clustering of Weibel-Palade bodies in endothelial cells. Arteriosclerosis, thrombosis, and vascular biology. Jan 2006;26(1):49–55.

4. Seidizadeh O, Eikenboom JCJ, Denis CV, et al. von Willebrand disease. Nature Reviews Disease Primers. 2024/07/25 2024;10(1):51. doi:10.1038/s41572-024-00536-8

5. Atiq F, O’Donnell JS. Novel functions for von Willebrand factor. Blood. 2024/09/19/ 2024;144(12):1247–1256. doi:10.1182/blood.2023021915

6. Zuchtriegel G, Uhl B, Puhr-Westerheide D, et al. Platelets Guide Leukocytes to Their Sites of Extravasation. PLoS Biol. May 2016;14(5):e1002459. doi:10.1371/journal.pbio.1002459

7. Braun LJ, Stegmeyer RI, Schäfer K, et al. Platelets docking to VWF prevent leaks during leukocyte extravasation by stimulating Tie-2. Blood. Jul 30 2020;136(5):627–639. doi:10.1182/blood.2019003442

8. Jevon D, Deng K, Hallahan N, et al. Local activation of focal adhesion kinase orchestrates the positioning of presynaptic scaffold proteins and Ca(2+) signalling to control glucose-dependent insulin secretion. Elife. May 13 2022;11 doi:10.7554/eLife.76262

9. Lopes da Silva M, Cutler DF. von Willebrand factor multimerization and the polarity of secretory pathways in endothelial cells. Blood. Jul 14 2016;128(2):277–85. doi:10.1182/blood-2015-10-677054

10. Weiss HJ, Sussman, II, Hoyer LW. Stabilization of factor VIII in plasma by the von Willebrand factor. Studies on posttransfusion and dissociated factor VIII and in patients with von Willebrand’s disease. J Clin Invest. Aug 1977;60(2):390–404. doi:10.1172/jci108788

11. Nightingale TD, White IJ, Doyle EL, et al. Actomyosin II contractility expels von Willebrand factor from Weibel–Palade bodies during exocytosis. Journal of Cell Biology. 2011;194(4):613–629. doi:10.1083/jcb.201011119

12. Nightingale TD, Pattni K, Hume AN, Seabra MC, Cutler DF. Rab27a and MyRIP regulate the amount and multimeric state of VWF released from endothelial cells. Blood. 2009;113(20):5010–5018. doi:10.1182/blood-2008-09-181206

13. Stehbens SJ, Paszek M, Pemble H, Ettinger A, Gierke S, Wittmann T. CLASPs link focal-adhesion-associated microtubule capture to localized exocytosis and adhesion site turnover. Nature cell biology. Jun 2014;16(6):561–73. doi:10.1038/ncb2975

14. Bjørnestad SA, Guadagno NA, Kjos I, Progida C. Rab33b-exocyst interaction mediates localized secretion for focal adhesion turnover and cell migration. iScience. 2022/05/20/ 2022;25(5):104250. doi:10.1016/j.isci.2022.104250

15. Fourriere L, Kasri A, Gareil N, et al. RAB6 and microtubules restrict protein secretion to focal adhesions. Journal of Cell Biology. 2019;218(7):2215–2231. doi:10.1083/jcb.201805002

16. Han X, Li P, Yang Z, et al. Zyxin regulates endothelial von Willebrand factor secretion by reorganizing actin filaments around exocytic granules. Nature Communications. 2017/03/03 2017;8(1):14639. doi:10.1038/ncomms14639

17. El-Mansi S, Robinson CL, Kostelnik KB, et al. Proximity proteomics identifies septins and PAK2 as decisive regulators of actomyosin-mediated expulsion of von Willebrand factor. Blood. 2023;141(8):930–944. doi:10.1182/blood.2022017419

18. El-Mansi S, Mitchell TP, Mobayen G, et al. Myosin-1C augments endothelial secretion of von Willebrand factor by linking contractile actomyosin machinery to the plasma membrane. Blood Advances. 2024;8(17):4714–4726. doi:10.1182/bloodadvances.2024012590

19. Nightingale TD, McCormack JJ, Grimes W, et al. Tuning the endothelial response: differential release of exocytic cargos from Weibel-Palade bodies. Journal of thrombosis and haemostasis : JTH. Sep 2018;16(9):1873–1886. doi:10.1111/jth.14218

20. Michaux G, Abbitt KB, Collinson LM, Haberichter SL, Norman KE, Cutler DF. The physiological function of von Willebrand’s factor depends on its tubular storage in endothelial Weibel-Palade bodies. Developmental cell. Feb 2006;10(2):223–32. doi:10.1016/j.devcel.2005.12.012

21. Stevenson NL, White IJ, McCormack JJ, Robinson C, Cutler DF, Nightingale TD. Clathrin-mediated post-fusion membrane retrieval influences the exocytic mode of endothelial Weibel-Palade bodies. Journal of cell science. Aug 1 2017;130(15):2591–2605. doi:10.1242/jcs.200840

22. Nightingale TD, White IJ, Doyle EL, et al. Actomyosin II contractility expels von Willebrand factor from Weibel-Palade bodies during exocytosis. The Journal of cell biology. Aug 22 2011;194(4):613–29. doi:jcb.201011119 [pii]10.1083/jcb.201011119

23. El-Mansi S, Robinson CL, Kostelnik KB, et al. Proximity proteomics identifies septin and PAK2 as decisive regulators of actomyosin expulsion of von Willebrand factor. Blood. 2022;doi:10.1182/blood.2022017419

24. Mehta V, Pang KL, Rozbesky D, et al. The guidance receptor plexin D1 is a mechanosensor in endothelial cells. Nature. Feb 2020;578(7794):290–295. doi:10.1038/s41586-020-1979-4

25. van Geemen D, Smeets MWJ, van Stalborch A-MD, et al. F-Actin–Anchored Focal Adhesions Distinguish Endothelial Phenotypes of Human Arteries and Veins. Arteriosclerosis, Thrombosis, and Vascular Biology. 2014;34(9):2059–2067. doi:doi:10.1161/ATVBAHA.114.304180

26. Collins C, Osborne LD, Guilluy C, et al. Haemodynamic and extracellular matrix cues regulate the mechanical phenotype and stiffness of aortic endothelial cells. Nature Communications. 2014/06/11 2014;5(1):3984. doi:10.1038/ncomms4984

27. Zhao ZS, Manser E, Loo TH, Lim L. Coupling of PAK-interacting exchange factor PIX to GIT1 promotes focal complex disassembly. Mol Cell Biol. Sep 2000;20(17):6354–63. doi:10.1128/mcb.20.17.6354-6363.2000

28. Audebert S, Navarro C, Nourry C, et al. Mammalian Scribble forms a tight complex with the betaPIX exchange factor. Curr Biol. Jun 8 2004;14(11):987–95. doi:10.1016/j.cub.2004.05.051

29. Davidson A, Tyler J, Hume P, Singh V, Koronakis V. A kinase-independent function of PAK is crucial for pathogen-mediated actin remodelling. PLoS Pathog. Aug 2021;17(8):e1009902. doi:10.1371/journal.ppat.1009902

30. Liaqat I, Hilska I, Saario M, et al. Spatially targeted chemokine exocytosis guides transmigration at lymphatic endothelial multicellular junctions. The EMBO Journal. 2024;43(15):3141–3174. doi:10.1038/s44318-024-00129-x

31. Gan WJ, Do OH, Cottle L, et al. Local Integrin Activation in Pancreatic β Cells Targets Insulin Secretion to the Vasculature. Cell Rep. Sep 11 2018;24(11):2819-2826.e3. doi:10.1016/j.celrep.2018.08.035

32. van Geemen D, Smeets MW, van Stalborch AM, et al. F-actin-anchored focal adhesions distinguish endothelial phenotypes of human arteries and veins. Arteriosclerosis, thrombosis, and vascular biology. Sep 2014;34(9):2059–67. doi:10.1161/ATVBAHA.114.304180

33. Zenner HL, Collinson LM, Michaux Gg, Cutler DF. High-pressure freezing provides insights into Weibel-Palade body biogenesis. Journal of Cell Science. 2007;120(12):2117–2125. doi:10.1242/jcs.007781

34. Nass J, Koerdt SN, Biesemann A, et al. Tip-end fusion of a rod-shaped secretory organelle. Cell Mol Life Sci. Jun 4 2022;79(6):344. doi:10.1007/s00018-022-04367-2

35. Steffes LC, Cheng P, Quertermous T, Kumar ME. von Willebrand Factor Is Produced Exclusively by Endothelium, Not Neointima, in Occlusive Vascular Lesions in Both Pulmonary Hypertension and Atherosclerosis. Circulation. Aug 2 2022;146(5):429–431. doi:10.1161/circulationaha.121.058427

36. Lagrange J, Worou ME, Michel JB, et al. The VWF/LRP4/αVβ3-axis represents a novel pathway regulating proliferation of human vascular smooth muscle cells. Cardiovasc Res. Jan 29 2022;118(2):622–637. doi:10.1093/cvr/cvab042

37. Farrugia AJ, Calvo F. Cdc42 regulates Cdc42EP3 function in cancer-associated fibroblasts. Small GTPases. 2017/01/02 2017;8(1):49–57. doi:10.1080/21541248.2016.1194952

38. Joberty G, Perlungher RR, Sheffield PJ, et al. Borg proteins control septin organization and are negatively regulated by Cdc42. Nature Cell Biology. 2001/10/01 2001;3(10):861–866. doi:10.1038/ncb1001-861

39. Zhu J, Zhou Q, Xia Y, et al. GIT/PIX Condensates Are Modular and Ideal for Distinct Compartmentalized Cell Signaling. Molecular Cell. 2020/09/03/ 2020;79(5):782-796.e6. doi:10.1016/j.molcel.2020.07.004

40. Hiroyasu S, Stimac GP, Hopkinson SB, Jones JCR. Loss of β-PIX inhibits focal adhesion disassembly and promotes keratinocyte motility via myosin light chain activation. Journal of Cell Science. 2017;130(14):2329–2343. doi:10.1242/jcs.196147

41. Manneville J-B, Etienne-Manneville S, Skehel P, Carter T, Ogden D, Ferenczi M. Interaction of the actin cytoskeleton with microtubules regulates secretory organelle movement near the plasma membrane in human endothelial cells. Journal of Cell Science. 2003;116(19):3927–3938. doi:10.1242/jcs.00672

42. Noordstra I, van den Berg CM, Boot FWJ, et al. Organization and dynamics of the cortical complexes controlling insulin secretion in β-cells. Journal of Cell Science. 2022;135(3)doi:10.1242/jcs.259430

43. Miteva KT, Pedicini L, Wilson LA, et al. Rab46 integrates Ca(2+) and histamine signaling to regulate selective cargo release from Weibel-Palade bodies. The Journal of cell biology. Jul 1 2019;218(7):2232–2246. doi:10.1083/jcb.201810118

44. Xu L, Qiu Y, Li Y, Wei Y, Wan Y, Deng W. Tissue dynamics of von Willebrand factor characterized by a novel fluorescent protein–von Willebrand factor chimera. Journal of Thrombosis and Haemostasis. 2022;20(1):208–221. doi:10.1111/jth.15542

