## Supplemental Files for "Endothelial β-PIX (ARHGEF7) drives exocytosis through enabling the dynamic reorganisation of the cytoskeleton"

**This PDF file includes:**

Table S1 and S2
Figs. S1-8

**Other Supplementary Materials for this manuscript include the following:**

Figure Legends for Movies

**Movie 1: Live cell imaging of FA-associated WPB exocytosis**. Confocal live cell imaging of HUVEC expressing GFP-VWF following PMA stimulation. GFP-VWF positive WPB (circled) can be seen to traffic toward the FA before exocytosis (indicated by arrows).

**Movie 2: "Hot-spots" of VWF secretion at the basement membrane**. WPB positioned below the nucleus fuse with the basement membrane where secreted VWF can be seen immobilised. Confocal live cell imaging of HUVEC expressing GFP-VWF following PMA stimulation. Region of interest covers the nucleus. Movies are processes with Z-depth colour coding whereby red>green>blue signifies the bottom, centre and top of the Z stack respectively. HUVEC expressing GFP-VWF. PMA stimulation (100ng/mL).

**Movie 3: Example of apical WPB exocytosis in the cell periphery.** GFP-VWF signal from WPBs fusing with the apical membrane is highly mobile and rapidly dissipates. Movies are processes with Z-depth colour coding whereby red>green>blue signifies the bottom, centre and top of the Z stack respectively. HUVEC expressing GFP-VWF. PMA stimulation (100ng/mL).

| Target name | Host species | Cat number | Supplier | Dilution | Method |
|---|---|---|---|---|---|
| **VWF** | Rabbit | A0082 | Dako | 1 in 10000 | IF |
| **VWF** | Sheep | AHP062 | Bio-Rad | 1 in 10000 | IF |
| **Vinculin** | Mouse | 14-9777-80 | eBioscience ™ | 1 in 200 | IF |
| **ARHGEF7/B-PIX** | Mouse | sc-393184 | Santa Cruz | 1 in 200 | IF |
| **GIT1** | Rabbit | 26247-1-AP | Protein Tech | 1 in 200 / 1 in 1000 | IF and WB |
| **GIT2** | Rabbit | GTX133285-S | Gene Tex | 1 in 500 | WB |
| **Scribble** | Rabbt | 27083-1-AP | Protein Tech | 1 in 1000 | WB |
| **Septin 7** | Rabbit | 18911 | Tecan | 1 in 1000 | IF |
| **NMIIA** | Rabbit | EPR8965 | Abcam | 1 in 200 | IF |
| **TUBULIN** | Mouse | T5201 | Sigma Aldrich | 1 in 3000 | WB |
| **GAPDH** | Mouse | 60004-1-Ig | Protein Tech | 1 in 3000 | WB |
| **GAPDH** | Rabbit | 10494-1-AP | Protein Tech | 1 in 3000 | WB |
| **MLC2 sampler kit** | Rabbit | 9776 | Cell Signalling | 1 in 200 / 1 in 1000 | IF and WB |
| **Actin Reorganization Antibody Sampler Kit** | Rabbit | 9967 | Cell Signalling | 1 in 1000 | WB |
| **Total PAK2** | Rabbit | 2608 | Cell Signalling | 1 in 1000 | WB |
| **PPAK2 s20** | Rabbit | 2607 | Cell Signalling | 1 in 1000 | WB |
| **PPAK2 thr402** | Rabbit | 2601 | Cell Signalling | 1 in 1000 | WB |
| **PAK2** | Mouse | MA5-15527 | Thermo Fisher Scientific, | 1 in 1000 | WB |
| **anti Rabbit IgG NIR 800** | Donkey | A21057 | Licor | 1 in 15000 | WB |
| **Anti-Mouse IgG NIR 680** | Goat | 92668071 | Licor | 1 in 15000 | WB |
| **GFP** | Rat | 3H9 | Chromotek | 1 in 1000 | WB |

Table S1: Details of antibodies used in the investigation.

| Construct | Source | Citation |
|---|---|---|
| **LifeAct-GFP** | A kind gift from the Baum Laboratory. | Riedl J, Crevenna AH, Kessenbrock K, et al. Lifeact: a versatile marker to visualize F-actin. Nat Methods. 2008;5(7):605-607. |
| **P.sel.lum.mCherry** | Our laboratory | Nightingale TD, White IJ, Doyle EL, et al. Actomyosin II contractility expels von Willebrand factor from Weibel-Palade bodies during exocytosis. J Cell Biol. 2011;194(4):613-629 |
| **Vinculin-mCherry** | pmCherry-Vinculin was a gift from Chinten James Lim (Addgene plasmid # 80024 ; http://n2t.net/addgene:80024 ; RRID:Addgene_80024) | Two modes of integrin activation form a binary molecular switch in adhesion maturation. Lee HS, Anekal P, Lim CJ, Liu CC, Ginsberg MH. Mol Biol Cell. 2013 May;24(9):1354-62. doi: 10.1091/mbc.E12-09-0695. Epub 2013 Mar 6. 10.1091/mbc.E12-09-0695 PubMed 23468527 |
| **GFP-VWF** | a gift from J. Voorberg and J.A. Van Mourik (Sanquin Research Laboratory, Amsterdam, The Netherlands). | Romani de Wit T, Rondaij MG, Hordijk PL, Voorberg J, van Mourik JA. Real-time imaging of the dynamics and secretory behavior of Weibel-Palade bodies. Arterioscler Thromb Vasc Biol. 2003;23(5):755-761. |
| **NMIIA-GFP** | A kind gift from the Baum Laboratory. | Curran S, Strandkvist C, Bathmann J, de Gennes M, Kabla A, Salbreux G, Baum B. Myosin II Controls Junction Fluctuations to Guide Epithelial Tissue Ordering. Dev Cell. 2017 Nov 20;43(4):480-492.e6. doi: 10.1016/j.devcel.2017.09.018. Epub 2017 Oct 26. PMID: 29107560; PMCID: PMC5703647. |
| **Cdc42-GFP** | A kind gift from Yuki Goda. | |
| **Cdc42-GFP(N17)** | A kind gift from Yuki Goda. | |
| **Cdc42-GFP(V12)** | A kind gift from Yuki Goda. | |
| **ARHGEF7-GFP constructs** | A kind gift from Anthony Davidson and Vassilis Koronakis | Davidson A, Tyler J, Hume P, Singh V, Koronakis V. A kinase-independent function of PAK is crucial for pathogen-mediated actin remodelling. PLoS Pathog. 2021 Aug 30;17(8):e1009902. doi: 10.1371/journal.ppat.1009902. PMID: 34460869; PMCID: PMC8432889. |

Table S2: Source of plasmid DNA constructs used for the transfection of HUVEC.

Figure S1

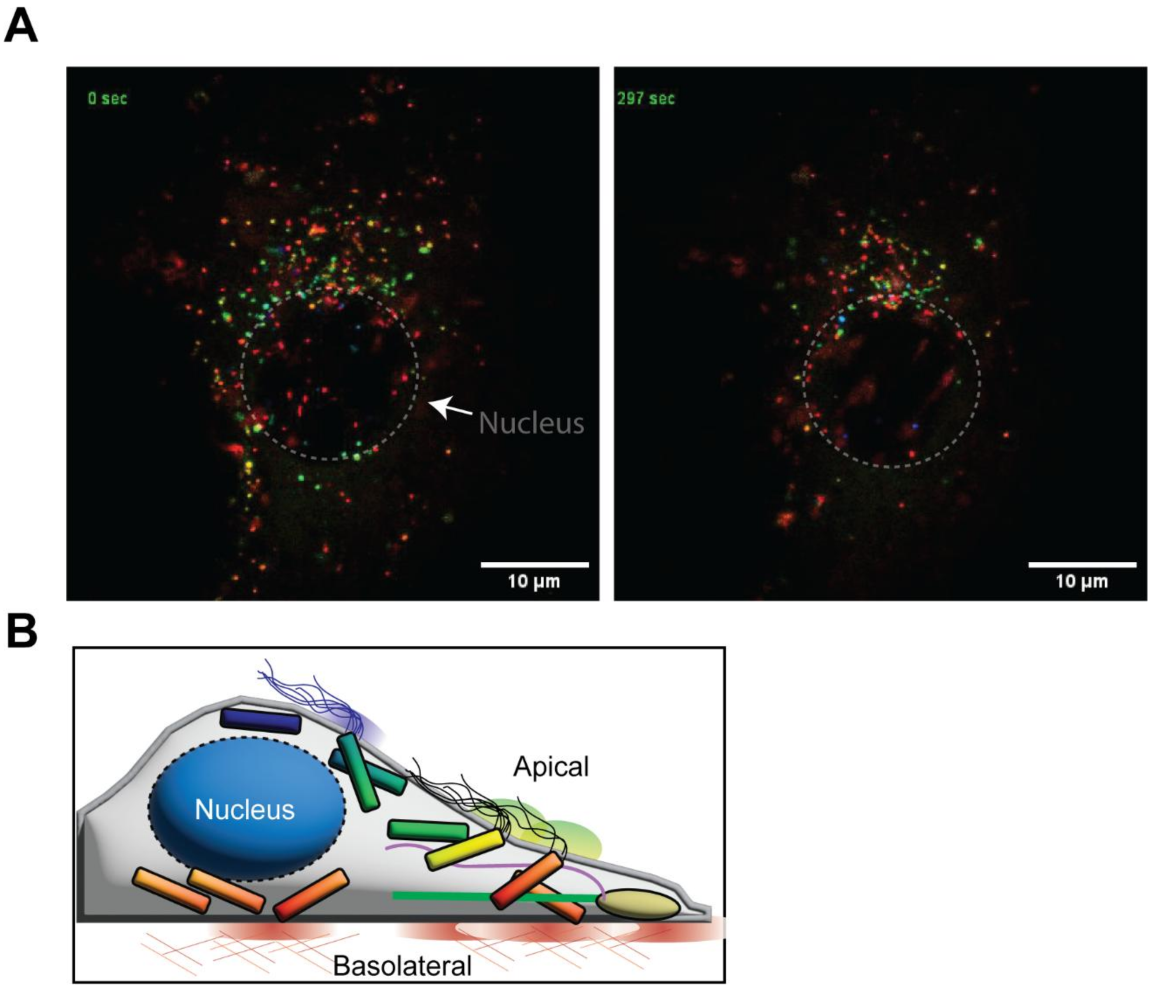


**Figure S1.** (A) Confocal images and (B) Schematic of live cell imaging approach to study polarity of WPB exocytosis.

Figure S2

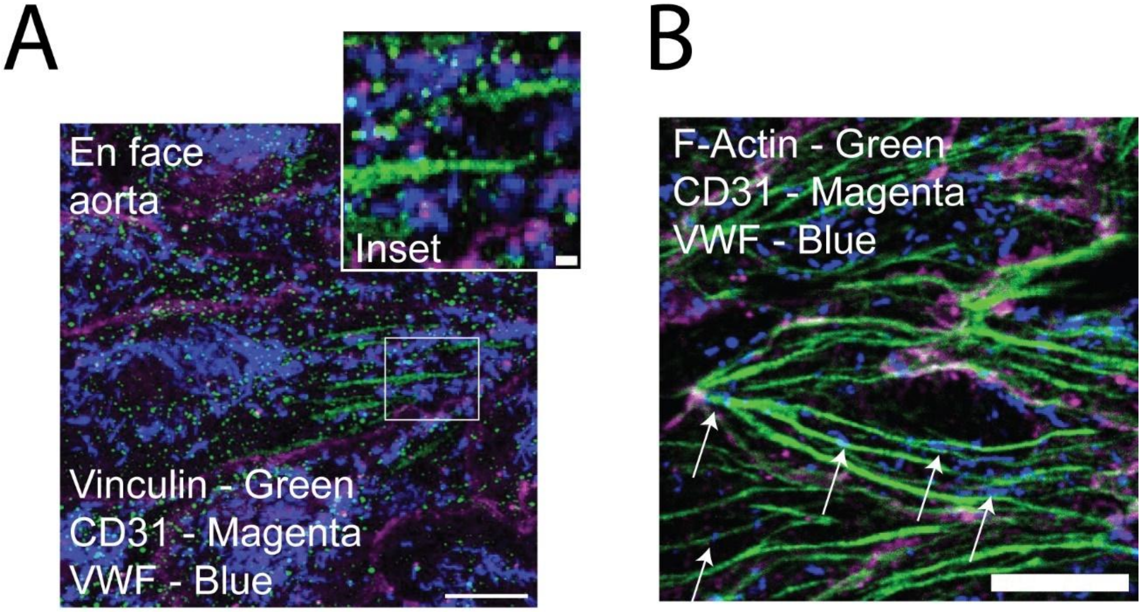


**Figure S2**: Localisation of Weibel-Palade bodies in murine aortic endothelial cells in vivo (A) *En face* confocal imaging of murine aortae indicated the WPBs (VWF-Blue) can be observed juxtaposed to focal adhesions (Vinculin – Green, PECAM-1 – Magenta) and (B) are also distributed across transverse F-actin stress fibres (Phalloidin-568- Green). Scale bar 10 µm.

Figure S3

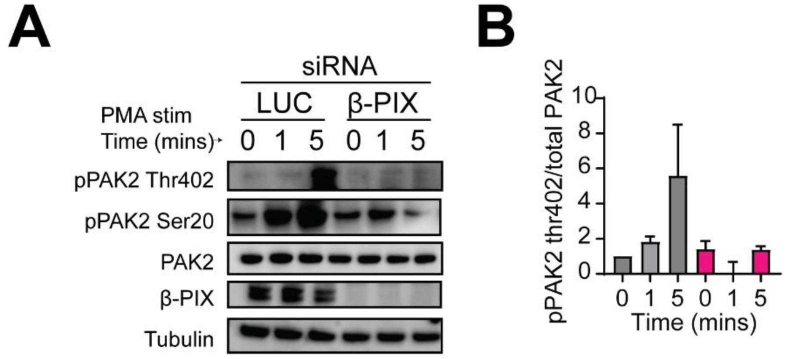


**Figure S3.** **(A)** PAK2 activation β-PIX depleted cells were determined through immunoblotting of total PAK2 and phospho-PAK2 (Thr402 and Ser20). **(B)** Densitometric analysis determined a time dependent increase in phospho-PAK2 (Thr402) in response to PMA stimulation (100 ng/mL).

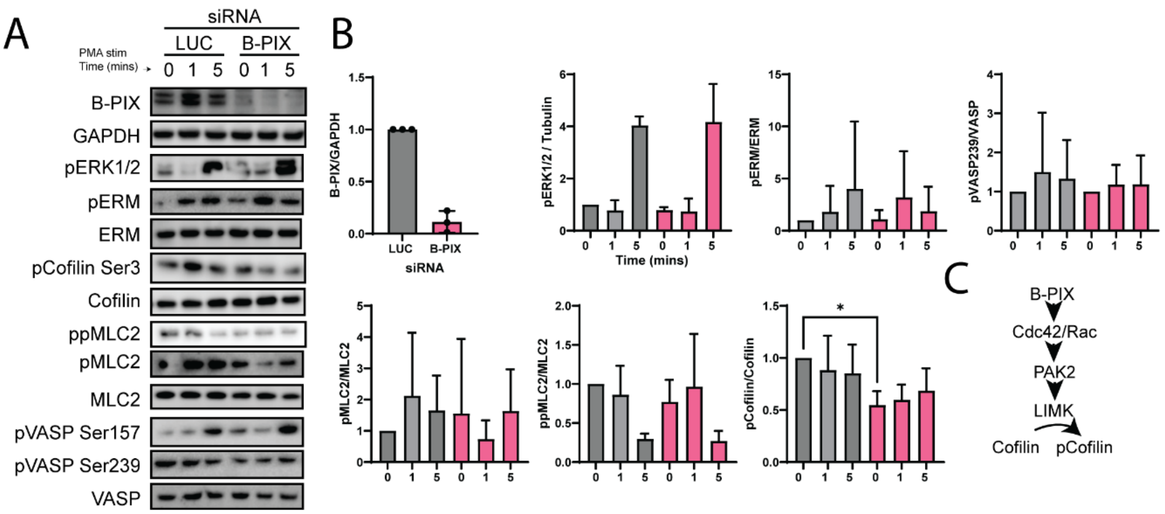


**Figure S4:** **(A)** Western immunoblotting and **(B)** densitometry to probe the relative phosphorylation of cytoskeletal remodelling proteins in control and β-PIX depleted cells under unstimulated and PMA stimulated conditions. N=3. *p<0.05 Ratio paired *t* test. **(C)** Schematic showing how β-PIX can lead to activation of PAK2 and regulation of cofilin.

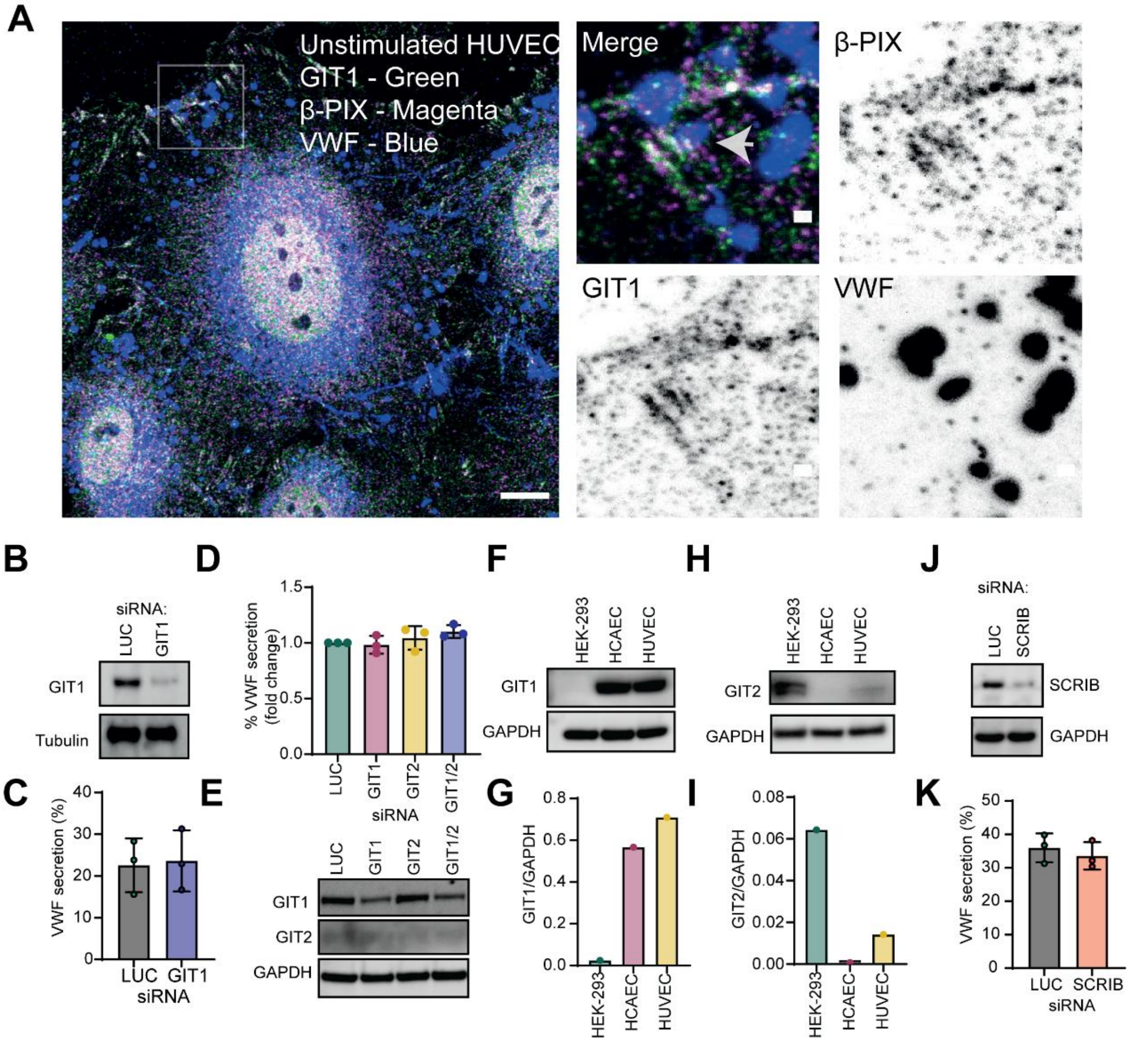


**Figure S5 (A)** Immunofluorescence of GIT1 (green), β-PIX (magenta) and VWF (blue) demonstrated an extent of colocalization at FAs and in the nucleus. Note the presence of WPBs (VWF-Blue) at Focal Adhesions. Scale bar 10 µm. Inset scale bar 1 µm **(B)** Transfection with 500 pM GIT1 siRNA reduced endogenous levels of GIT1 but did not affect **(C)** VWF secretion in response to PMA n=3. LUC = Luciferase **(D)** HUVEC were transfected with siRNA targeting LUC, GIT, GIT2 or a combination of GIT1 and GIT2. Neither siRNA effected VWF secretion, alone or in combination n=3. **(E)** Western blotting of GAPDH, GIT1 and GIT2 in lysates from siRNA transfected HUVEC. **(F-I)** Western blotting and densitometry of **(F&G)** GIT1 and **(H&I)** GIT2 protein levels in HEK-293, HCAEC and HUVEC lysates. **(J)** Immunoblotting of lysates from LUC and Scribble KD HUVEC. **(K)** Scribble deletion did not affect VWF secretion in response to PMA. N=3 *n.s* not statistically significant. *p<0.05 ****p<0.0001. Students *t* test.

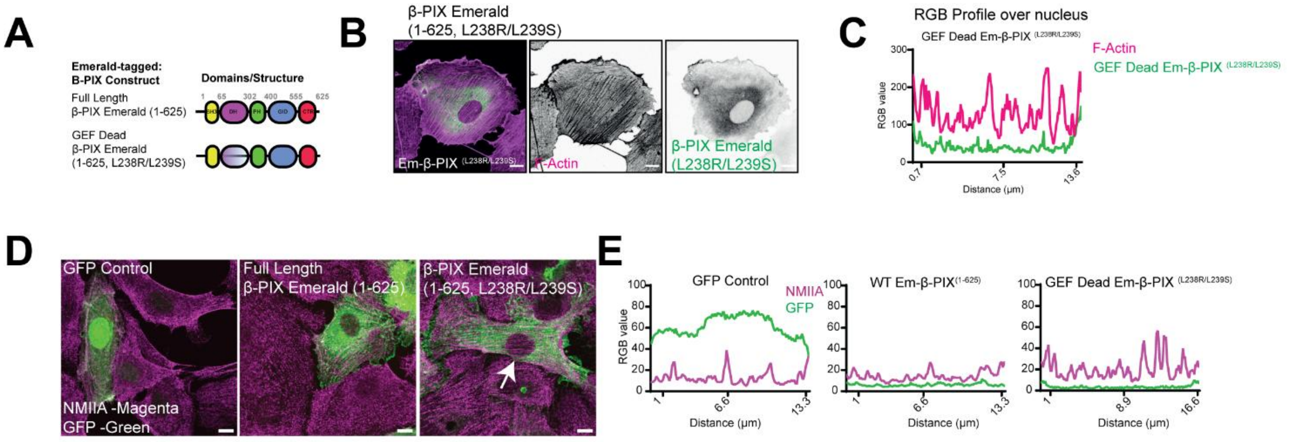


**Figure S6.** **(A)** Schematic representation of the domains and structure of Em-β-PIX$^{(1-165)}$ and Em-β-PIX$^{(L238R/L293S)}$ (GEF dead). **(B)** Confocal imaging and analysis of F-actin architecture determined that Em-β-PIX$^{(L238R/L293S)}$ expression phenocopied the effect of SH3 and SH3/DH truncated mutants **(C)** RGB profile plot of Em-β-PIX and F-Actin above the nucleus. **(D)** Analysis of the subcellular localisation of endogenous NMII demonstrated its misplacement above the nucleus on filamentous structures in HUVEC expressing the Em-β-PIX$^{(L238R/L293S)}$. Scale bars are 10 μm. **(E)** RGB profile plots of NMIIA and GFP/Em-β-PIX above the nucleus.

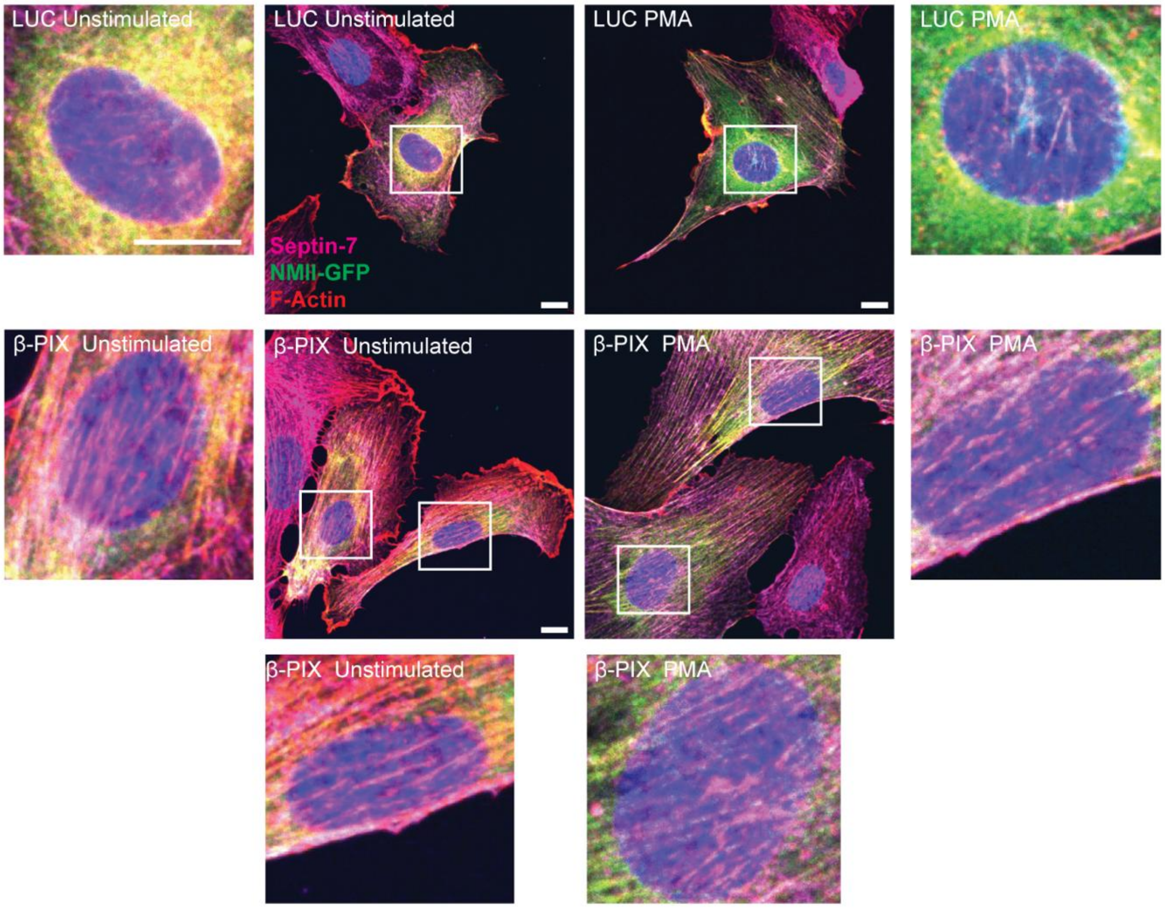


**Figure S7. β-PIX depletion results in prominent F-actin stress fibres above the nucleus which are decorated in Septin-7 and NMIIA-GFP.** Spinning disk microscopy of HUVEC under unstimulated or stimulated conditions. NMII-GFP (green), Septin-7 (Magenta) and F-actin (red)in control and β-PIX depletion HUVEC. Scale bar 10μm.

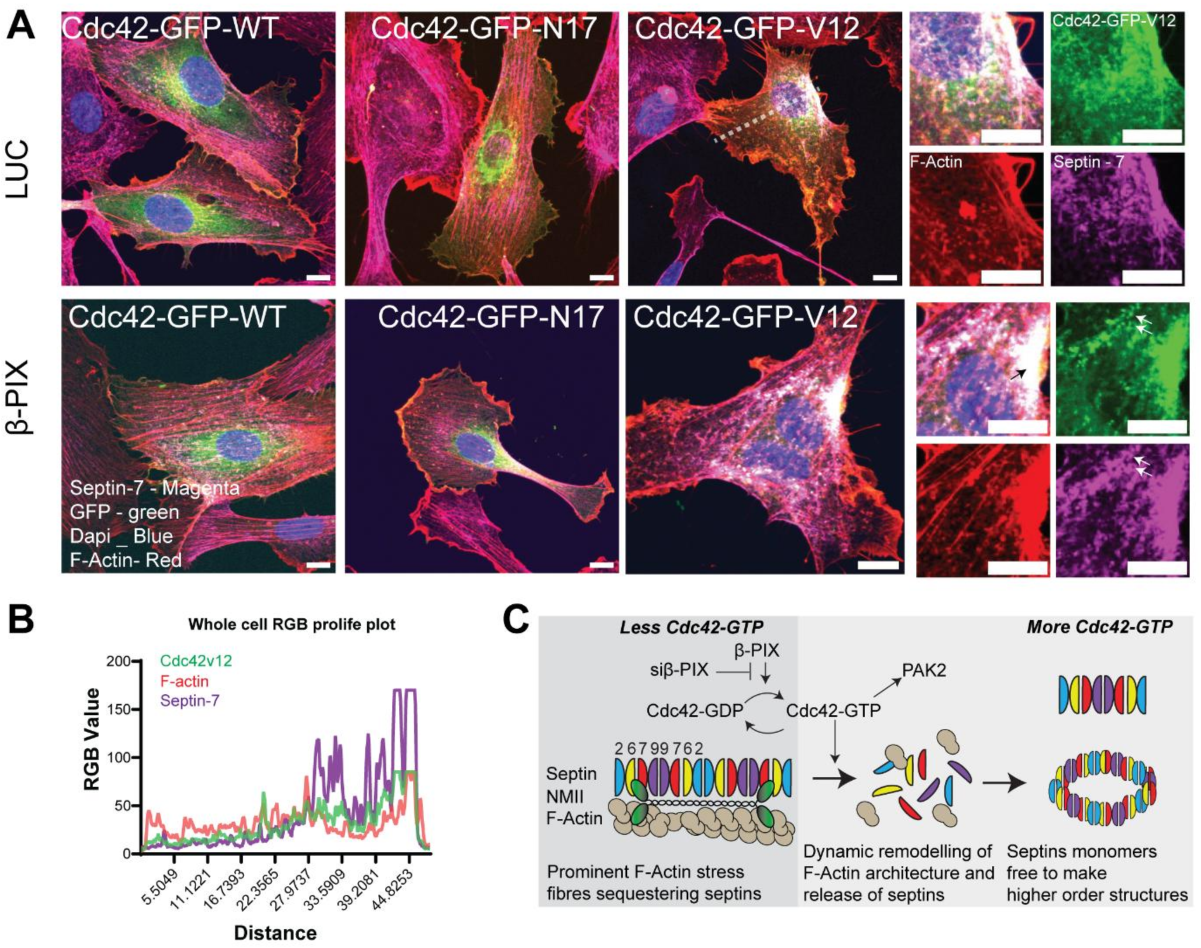


**Figure S8.** (A) Overexpression of a GFP-tagged, constitutively active Cdc42 (V12) construct demonstrated its colocalization with septin-7 (magenta) and actin (red). Cdc42-GFP (V12) rescued the cytoskeletal phenotype observed in β-PIX KD cells. Insets and arrows show the presence of septin-7, cdc42-GFP V12 and actin positive rings and filaments. (B) RGB profile plots across the whole cell shows the overlapping peaks of septins and cdc42V12. (C) Working schematic of a potential mechanism by which perturbing Cdc42-GDP>GTP cycling through B-PIX depletion may effect septin and actin dynamics.
